# TSpred: a robust prediction framework for TCR-epitope interactions based on an ensemble deep learning approach using paired chain TCR sequence data

**DOI:** 10.1101/2023.12.04.570002

**Authors:** Ha Young Kim, Sungsik Kim, Woong-Yang Park, Dongsup Kim

**Affiliations:** Department of Bio and Brain Engineering, Korea Advanced Institute of Science and Technology, Daejeon 34141, South Korea; GENINUS Inc., Seoul, South Korea; Samsung Genome Institute, Samsung Medical Center, Seoul, South Korea; Department of Molecular Cell Biology, Sungkyunkwan University School of Medicine, Suwon, South Korea

## Abstract

Prediction of T-cell receptor (TCR)-epitope interactions is important for many applications such as cancer immunotherapy. However, due to the scarcity of available data, it is known to be a challenging task particularly for novel epitopes. Here, we propose TSpred, a new ensemble deep learning approach for the pan-specific prediction of TCR binding specificity based on paired chain TCR data. This method combines the predictive power of CNN and the attention mechanism to capture the patterns underlying TCR-epitope interactions. In particular, we design a reciprocal attention mechanism which contributes to higher model generalizability to unseen epitopes. We perform a comprehensive evaluation of our model and observe that TSpred achieves state-of-the-art performances in both seen and unseen epitope specificity prediction tasks. Our model performs consistently well across both of the two widely used negative sampling strategies, while avoiding the potential bias associated with each strategy. Also, compared to other predictors, it is more robust to bias related to peptide imbalance in the dataset. In addition, the reciprocal attention component of our model allows for model interpretability by capturing structurally important binding regions. Results indicate that TSpred is a robust and reliable method for the task of TCR-epitope binding prediction.

## INTRODUCTION

T-cells are known to play a critical role in adaptive immune responses, detecting and eliminating infected cells in the body upon activation of T-cell receptors (TCRs). The TCRs are activated when they bind to a peptide that is presented on the surface of infected cells by the major histocompatibility complex (pMHC). TCR sequences have an enormously large sequence diversity, which enables the recognition of a large number of different epitopes, thereby protecting the host from a wide variety of pathogens (1). This sequence diversity is observed in the complementarity determining regions (CDRs) of the TCR. TCRs possess the property of cross-reactivity, such that the same TCR can bind to a number of different epitopes (2). At the same time, TCRs bind to epitopes in a highly specific manner, meaning that it is highly unlikely that a single TCR will bind to any randomly chosen epitope (3). Experimental approaches such as tetramer analysis (4) and single-cell TCR sequencing (5) are used to detect TCR-epitope interacting pairs. Despite an increasing amount of available data, it is still a challenge to predict which TCRs target specific epitopes. This is because the data available at the current moment is still too sparse compared to the huge sequence space of TCRs (6). In particular, the prediction of TCR binding specificities for unseen epitopes—epitopes not seen in the training data—is highly difficult, due to the lack of available data (7,8). Although many methods have been developed in this field, most of them fail to extrapolate well enough to unseen epitopes. Still, it is a very active area of research, as there are many practical applications associated with the prediction of TCR-epitope binding. For example, a reliable method for TCR-epitope binding prediction can be used for the prediction of immunogenic neoantigens, which has significant implications for cancer vaccine development (9).

Many machine learning and deep learning-based methods have been developed to predict the interaction between TCRs and epitopes. Deep learning-based methods use a wide variety of architectures, such as convolutional neural networks (CNNs) (10-13), long short-term memory (LSTM) (14), autoencoder (14), and attention mechanism (6,15,16). Some tools, such as TITAN (6), ImRex (13), pMTnet (17), epiTCR (18), and TEINet (19), consider only the information of the TCR beta chain. Other methods, such as ERGO (14), MixTCRpred (16), NetTCR-2.0 (12), NetTCR-2.1 (11), and NetTCR-2.2 (10), take into consideration the paired alpha and beta chain information. A recent benchmark study from the IMMREP 2022 workshop (20) has reported that using paired chain data leads to more accurate predictions. Furthermore, some works (10,11,16) make a distinction between epitope-specific and panspecific predictors. Epitope-specific predictors are specifically trained and tested for predicting the binding TCRs for the particular epitope, whereas pan-specific predictors can be applied to the prediction of binding TCRs for any given epitope. The authors of NetTCR-2.1 and MixTCRpred (11,16) pointed out that models tend to perform worse when trained in a pan-specific manner compared to an epitope-specific manner. However, it is important for a model to be a reliable pan-specific predictor so that it can generalize well to unseen epitopes. Also, a recent study (7) investigated how the peptide imbalance in the dataset affects the performances of different predictors. The authors found that the peptide imbalance leads to the overestimation of model performances and that the models learn only on a few number of peptides tha appear most frequently in the dataset.

In this work, we present TSpred (**T**-cell receptor binding **S**pecificity **pred**ictor), a panspecific approach for the prediction of TCR binding specificity using an ensemble of a CNN-based model and an attention-based model on paired chain TCR data. We leverage the predictive power of the CNN and the attention mechanism to build a robust model that can generalize well to unseen epitopes. In particular, we adopt a reciprocal attention mechanism which is specifically designed to capture the patterns underlying TCR-epitope binding. Based on a thorough evaluation of the model and comparison with other recent methods, we show that our model achieves state-of-the-art performances, in both seen epitope datasets and unseen epitope datasets. Also, based on an assessment of our model on a balanced dataset generated by down-sampling, we find that our model is the most robust to bias caused by peptide imbalance in the dataset. Furthermore, we analyze the attention maps generated by the reciprocal attention layer and show that our model can capture the structurally important residue pairs that contribute to TCR-epitope binding.

## MATERIAL AND METHODS

### Negative sampling strategies

Two approaches for sampling negatives are widely used in this field: random shuffling and sampling from negative control data (20). In the random shuffling approach, for each peptide-TCR pair, negative samples are generated by random sampling from TCRs binding to other peptides. A limitation of this strategy is that it must be conducted separately within the train and test partitions. Otherwise, a positive TCR in the training set can appear as a negative in the test set and vice versa, leading to a significant bias. This strategy can be difficult to use in cases where there are only a few peptides with an imbalanced number of samples within each partition. The second strategy is sampling from negative control TCR data without known epitope specificity, obtained from healthy individuals. A number of works (8,13,20,21) have pointed out the problem of this approach, which is the bias that arises due to the difference in the distributions of the positive and negative TCR sequences. In some cases, this makes the classification problem too easy, resulting in some predictive models achieving exceedingly high performances with ROC-AUC greater than 0.9 (8,18,22). In our study, we use both strategies depending on the type of datasets, while avoiding limitations and potential bias accompanying each strategy.

### Dataset

We conduct a comprehensive and rigorous evaluation of our model on four datasets, two of which are provided by the authors of NetTCR-2.2 (10), and two of which are newly constructed based on the previous two datasets (Table 1). The NetTCR_full dataset, refered to as the ‘full dataset’ in NetTCR-2.2 (10), is derived from public databases such as VDJDB (23) and IEDB (24), and a 10x sequencing study (25). The data is restricted to human and MHC class I data and contains paired chain information, including all three CDR sequences for both α and β chains. The positive pairs in this data, comprising 6353 examples across 26 peptides, are randomly split into five partitions. Within each partition, negative pairs are sampled in a 1:5 ratio by random shuffling. Whereas the nested five-fold cross validation has been performed in NetTCR-2.2 (10), we use a modified nested five-fold cross validation with only one inner loop and five outer loops to save computational time (Supplementary Fig S1). This training strategy is used throughout this work. In addition, we use the IMMREP dataset which has been generated from the IMMREP 2022 benchmark study (20) and post-processed by the authors of NetTCR-2.2 (10). This dataset is also a five-fold cross validation dataset with five randomly split partitions. The negative samples in this dataset have been generated by a combination of random shuffling and sampling from negative control data, with a final positive-to-negative ratio of 1:5. The negative control data come from the IMMREP benchmark study (20) and consist of 15,957 TCR sequences with no known binding specificity obtained by 10x sequencing from 11 control individuals. This dataset consists of 17 different peptides.

**Table 1.** Summary of the datasets used for model training and evaluation in this study.

| Dataset | NetTCR_full | IMMREP | NetTCR_bal | NetTCR_strict |
| --- | --- | --- | --- | --- |
| Task | Prediction for seen epitopes |  |  | Prediction for unseen epitopes |
| Training strategy | 5-fold cross validation (random split) |  |  | 5-fold cross validation (strict split) five times with five different random seeds |
| Data origin | NetTCR_full dataset from Jensen et al., 2023 (10) | IMMREP 2022 benchmark (20) | NetTCR_full | NetTCR_full |
| Negative sampling strategy | Random shuffling within each partition | Random shuffling (1:3) + negative controls (1:2) | Random shuffling within each partition | Negative controls |
| Number of peptides | 26 | 17 | 26 | 26 |
| Total data size (positive, negative) | (6353, 31368) | (1960, 9848) | (1893, 3786) | (6353, 6353) |
| Positive-to-negative ratio | 1:5 | 1:5 | 1:2 | 1:1 |

Based on NetTCR_full dataset, we create two other datasets, NetTCR_bal and NetTCR_strict. NetTCR_bal is a balanced dataset which is created in order to minimize the impact of peptide imbalance on the model performance. It is generated by combining all positive pairs in the NetTCR_full dataset and down-sampling the data to 100 samples for each peptide. Again, all the positive pairs are randomly split into five partitions, and negatives are generated in each partition by random shuffling in a 1:2 ratio, which is the limit of the positive-to-negative ratio that can be achieved with the reduced amount of data in each partition. Finally, NetTCR_strict dataset is constructed to assess our model on the task of predicting TCR specificity for unseen epitopes. When dividing all the positive data into five partitions, the ‘strict split’ method is used, so that peptides are non-overlapping across different partitions. In order to limit the influence of randomness, we split the data five times using five different random seeds, and conduct five-fold cross validation for each data split. For the negative samples, we randomly sample from the previously mentioned negative control data from IMMREP benchmark (20), with a positive-to-negative ratio of 1:1. We use negative controls here because each partition generated by the strict split contains such a small number of peptides with an imbalanced number of samples that it is not possible to achieve a 1:1 ratio by random shuffling in certain cases.

### Model Architecture

In this study, we propose an ensemble model of two different models, a CNN-based model and an attention-based model (Figure 1, Supplementary Note S1). In the CNN-based model (Figure 1a), each of the peptide and the 6 CDR sequences are one-hot encoded and forwarded through a convolution module. The convolution module is composed of a 1D convolutional layer with a kernel size of 2, a max pooling layer with a kernel size of 2, and a fully-connected layer. The outputs from each module are concatenated and passed through three fully-connected layers with a sigmoid activation to produce the final output.

**Figure 1.**
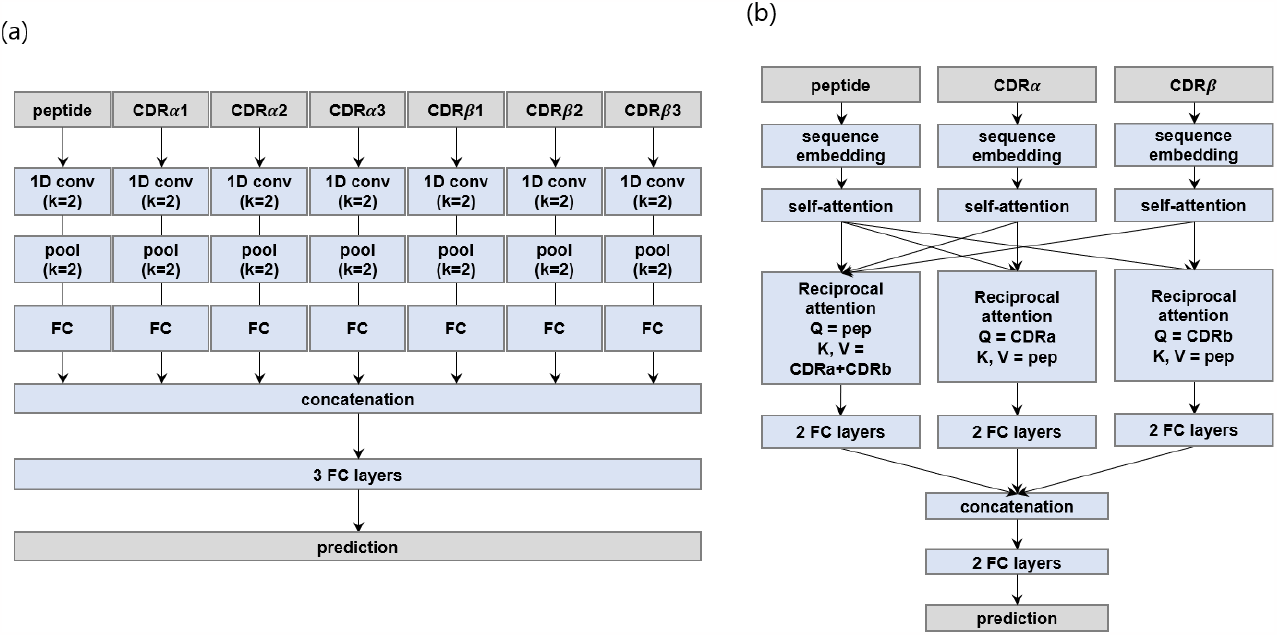
Overview of TSpred model architecture. (a) CNN-based model. Each of the peptide and 6 CDRs pass through a 1D convolutional layer, pooling layer, and a fully connected layer. All vectors are concatenated and forwarded through a series of fully connected layers to output the final value. (b) Attention-based model. In this model, the peptide, CDRα, and CDRβ sequences are the input. Each input is passed through a sequence embedding layer, self-attention layer, reciprocal attention layer, and two fully-connected layers. All vectors are concatenated and forwarded through a series of fully-connected layers to output the final value. The final TSpred model is an ensemble model of (a) and (b).

In the attention-based model (Figure 1b), the inputs are the peptide, CDRα, and CDRβ sequences, where the three CDRs from the α chain are concatenated and the three CDRs from the β chain are concatenated. Each input sequence is fed into a learnable embedding layer, followed by a multi-head self-attention layer. Then the vectors are each passed through a multi-head reciprocal attention layer, which takes the input sequence as the query and the other sequences as the key and value. Specifically, for the layer with the peptide as the query, the key and value are both the concatenated sequences of CDRα and CDRβ. For the layer with CDRα as the query, the key and value are both the peptide. The same applies to the layer with CDRβ as the query. This layer is designed so that the model can focus more on the relevant parts of the sequences of the interacting partners. After the reciprocal attention layer, each output is passed through two fully-connected layers. The three resulting vectors are then concatenated, flattened, and fed into two fully-connected layers with a sigmoid activation to produce the final output.

The final ensemble model of the CNN- and attention-based models is obtained by taking the average of the predictions of each model. We use different training hyperparameters for the CNN- and attention-based models (Supplementary Note S1). The criterion for choosing the number of epochs is based on the ROC-AUC on the validation set.

### Performance Evaluation

For the model performance evaluation, we report the average of the model performances on the test set across the five folds. We use the metrics of ROC-AUC (Area Under the Receiver Operating Characteristic Curve) and PR-AUC (Area Under the Precision-Recall Curve). We also report the accuracy, precision, specificity, recall and F1-score based on a cutoff of 0.5. These metrics are calculated as follows:

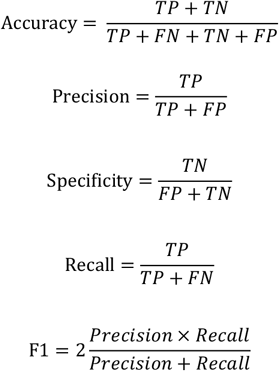

where TP, TN, FP, and FN are the number of True Positives, True Negatives, False Positives, and False Negatives, respectively.

### Comparison to Other Methods

We compare our method with four other recent state-of-the-art methods, TEINet (19), epiTCR (18), MixTCRpred (16), and NetTCR-2.2 (10), which use a variety of model architectures. TEINet and epiTCR take only the peptide and CDRβ3 as input, while MixTCRpred and NetTCR-2.2 take as input the peptide and all three CDRs from both α and β chains. TEINet is a method based on pre-trained encoders, and epiTCR is a method based on a random forest model. MixTCRpred is constructed using the transformer encoder architecture, and NetTCR-2.2 is built using CNNs. In this study, MixTCRpred is re-implemented based on the source code made available by the authors. For both MixTCRpred and NetTCR-2.2, the pan-specific model architectures are used for assessment. We compare all models using the same training, validation, and test datasets.

## RESULTS

### Prediction on Seen Epitopes

We evaluate the performances of the final TSpred model (TSpred_ensemble) as well as the individual model components (TSpred_CNN and TSpred_attention) and the other methods on NetTCR_full dataset using ROC-AUC and PR-AUC (Fig 2A, 2B). TSpred_CNN and TSpred_attention both achieve state-of-the-art results in terms of both metrics. By combining the predictive power of the two individual models, TSpred_ensemble achieves even higher performance, with a mean ROC-AUC of 0.86 and a mean PR-AUC of 0.66. For this dataset, the methods using the paired chain data outperform the methods using only beta chain data. Upon evaluation in terms of classification metrics, we observe performances similar to or better than the other methods in most cases (Supplementary Figure S2). We also inspect the performances of TSpred_ensemble in terms of ROC-AUC for each peptide in the dataset (Figure 3). The five most frequent peptides have an average ROC-AUC of 0.85, and the five least frequent peptides have an average ROC-AUC of 0.79. The peptide ELAGIGILTV, which has 426 positive samples, shows the highest performance with a ROC-AUC of 0.96. The peptide SLFNTVATLY, with 38 positive samples, achieves the lowest performance with a ROC-AUC of 0.65.

**Figure 2.**
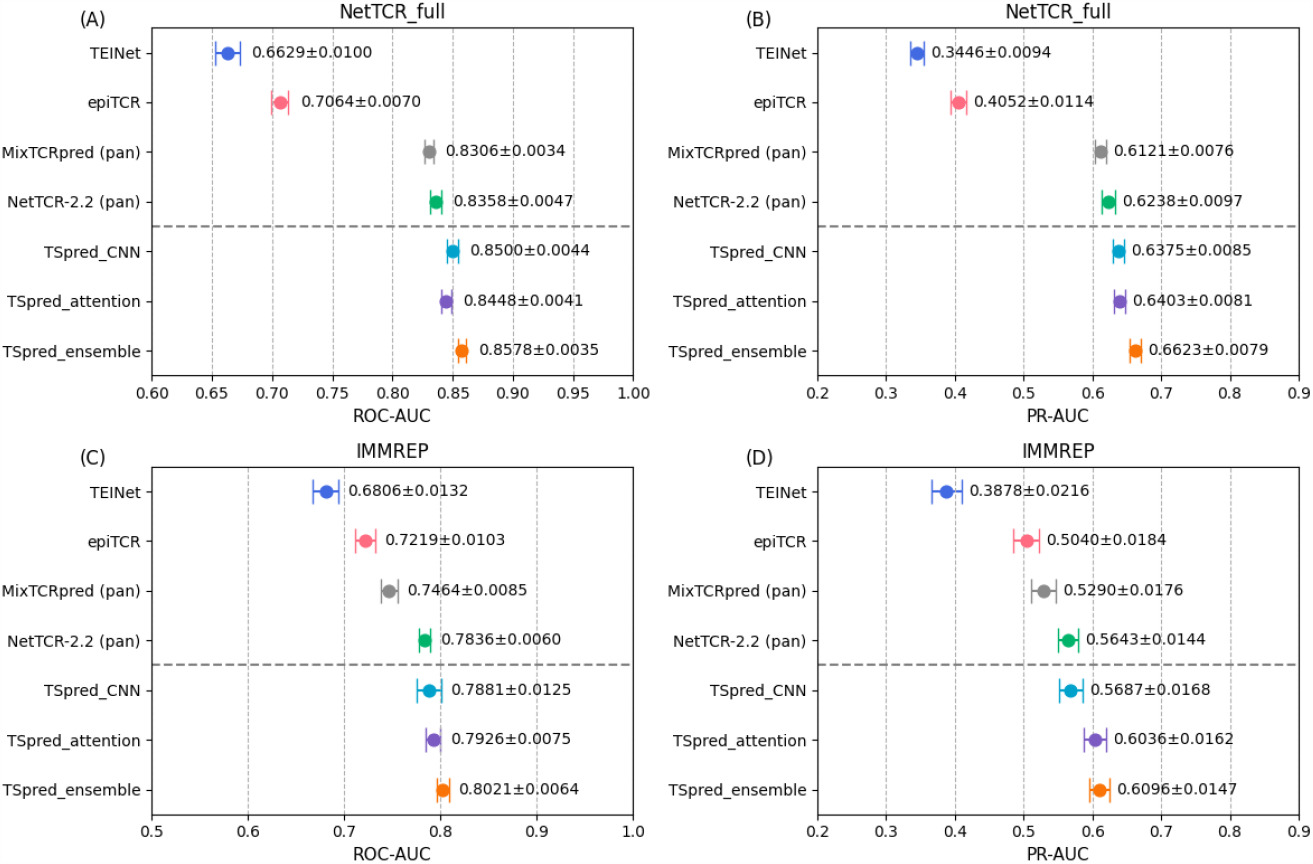
Performances of TSpred and other models on the seen epitope datasets. The colored dots represent the mean and the whiskers represent the standard deviation. (A) and (B) show the model performances in terms of ROC-AUC and PR-AUC on the NetTCR-full dataset, respectively. (C) and (D) show the model performances in terms of ROC-AUC and PR-AUC on the IMMREP dataset, respectively. The upper part of each subplot shows the results of the compared state-of-the-art methods, whereas the lower part of each subplot shows the results of the final TSpred model (TSpred_ensemble) and its individual components (TSpred_CNN and TSpred_attention).

**Figure 3.**
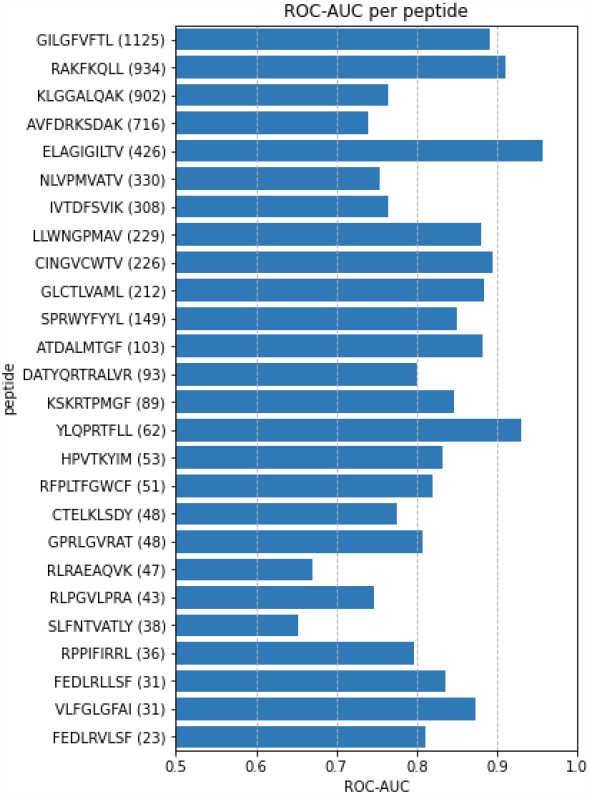
Performances of TSpred_ensemble in terms of ROC-AUC per peptide on the NetTCR_full dataset. The numbers shown in parentheses refer to the number of positive samples for each peptide.

Furthermore, we assess the performances of different methods on the IMMREP benchmark dataset for the task of predicting specificity for seen the epitopes (Fig 2C, 2D). Again, we observe that TSpred_CNN and TSpred_attention show better performances compared to the other tools, and that TSpred_ensemble even outperforms these two models. We once again observe that using the paired chain data as the model input leads to better prediction accuracy compared to using only beta chain data. In terms of classification metrics, our models show high accuracy, specificity, and F1-score (Supplementary Figure S3).

### Assessment on a Balanced Dataset

In order to rule out the bias caused by peptide imbalance on model performances, which has been pointed out by a recent study (7), we evaluate our model on a balanced dataset named NetTCR_bal, which is constructed by down-sampling the data from NetTCR_full dataset. We compare the performances of different methods in terms of ROC-AUC and PR-AUC (Figure 4). As expected, the performances drop significantly, due to the reduced amount of data.

**Figure 4.**
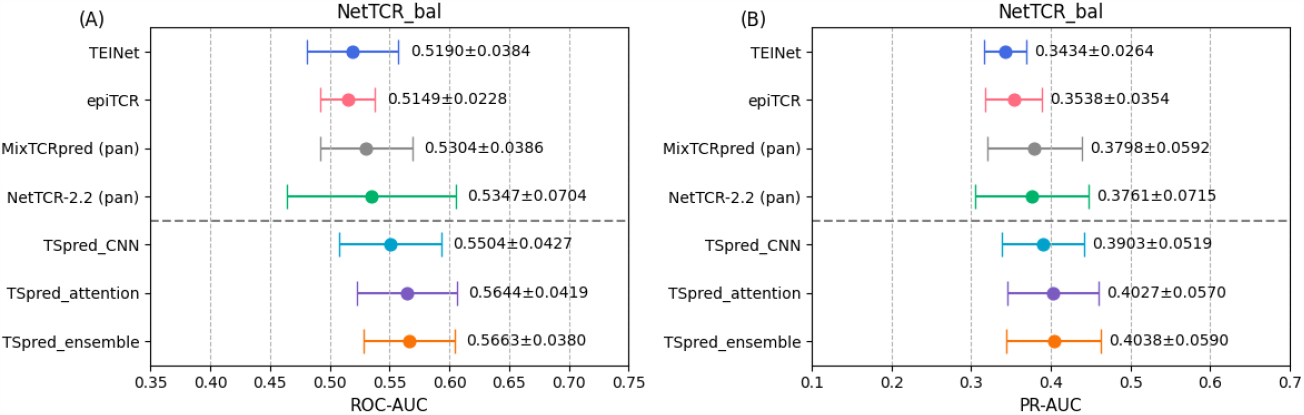
Performances of TSpred and other models on the NetTCR_bal dataset. The colored dots represent the mean and the whiskers represent the standard deviation. (A) and (B) show the model performances in terms of ROC-AUC and PR-AUC, respectively. The upper part of each subplot shows the results of the compared state-of-the-art methods, whereas the lower part of each subplot shows the results of the final TSpred model (TSpred_ensemble) and its individual components (TSpred_CNN and TSpred_attention).

Nevertheless, our models demonstrate higher performances compared to other methods. When evaluated using the classification metrics, our models show good accuracy, precision, and specificity (Supplementary Figure S4). Overall, these results demonstrate the robustness of TSpred, and show that TSpred is least influenced by the bias caused by the peptide imbalance in the dataset.

### Prediction on Unseen Epitopes

We next move onto the task of specificity prediction for unseen epitopes, which is a much more challenging problem. Model performances in terms of ROC-AUC and PR-AUC on NetTCR_strict dataset are analyzed (Figure 5). Although NetTCR_strict dataset contains negative samples derived from the negative control data, we observe that ROC-AUC values are mostly between 0.6 and 0.7, suggesting that the prediction task is not overly straightforward for the models. This implies that the negative control data used in this study is less biased than those of some of the previous studies (8,18,22) which lead to a very high ROC-AUC. We also notice that there is less variation among the results of different predictors, and that the predictors based on the beta chain data do not show results that are much different from the ones based on the paired chain data. We find that TSpred_ensemble achieves the highest performances among all methods in terms of ROC-AUC and PR-AUC. TSpred_attention demonstrates slightly better performances compared to TSpred_CNN, suggesting that the reciprocal attention mechanism helps the model to better capture the general features associated with TCR-epitope interactions. In terms of classification metrics, our methods demonstrate a good accuracy, precision, and specificity (Supplementary Figure S5). Noticeably, TEINet shows higher performances in the unseen epitope prediction task compared to the seen epitope prediction task. The authors of TEINet pre-trained their model on a large number of TCR sequences as well as a large number of epitope sequences (19). We speculate that this may have led to a higher generalizability of the model to unseen epitopes, which resulted in good predictive performances for this task.

**Figure 5.**
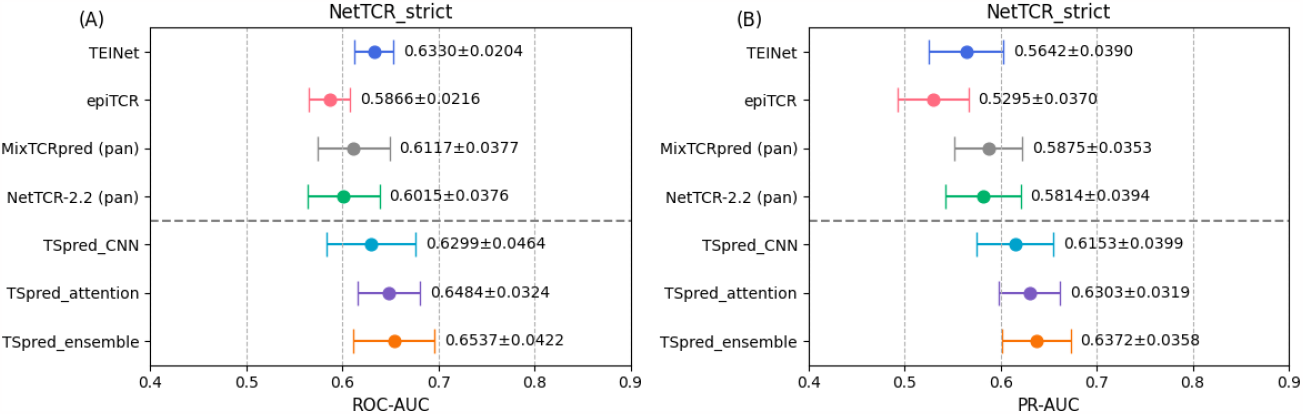
Performances of TSpred and other models on the NetTCR_strict dataset. The colored dots represent the mean and the whiskers represent the standard deviation. (A) and (B) show the model performances in terms of ROC-AUC and PR-AUC, respectively. The upper part of each subplot shows the results of the compared state-of-the-art methods, whereas the lower part of each subplot shows the results of the final TSpred model (TSpred_ensemble) and its individual components (TSpred_CNN and TSpred_attention).

### Attention Map Analysis

The reciprocal attention layer in TSpred_attention model has been conceived to capture the interaction patterns underpinning TCR-epitope binding. In order to examine whether our attention model can capture the structurally interacting residue pairs, we train and validate our model on the full NetTCR_full dataset and make predictions on a test dataset consisting of unseen peptide-TCR pairs derived from the STCRDab database (downloaded Aug. 2022) (26). We compare the experimentally determined 3D structures to the attention maps generated by the reciprocal attention layer (Figure 6). One example is the case with PDB code 4JFF, for which the attention map shows a high score for the L98 residue in CDRβ3 and the G6 residue in the peptide (Figure 6A). Upon examination of the structure, we find that the two residues are in close contact (3.1Å). Another example is the case with PDB code 3QEQ (Figure 6B). The attention map indicates a high score for the G97 residue in CDRβ3 and the T8 residue in the peptide. In the 3D structure, these two residues are in close contact with each other (4.8Å). These results indicate that the reciprocal attention layer used in our model can efficiently learn the patterns underlying the binding of TCRs and epitopes.

**Figure 6.**
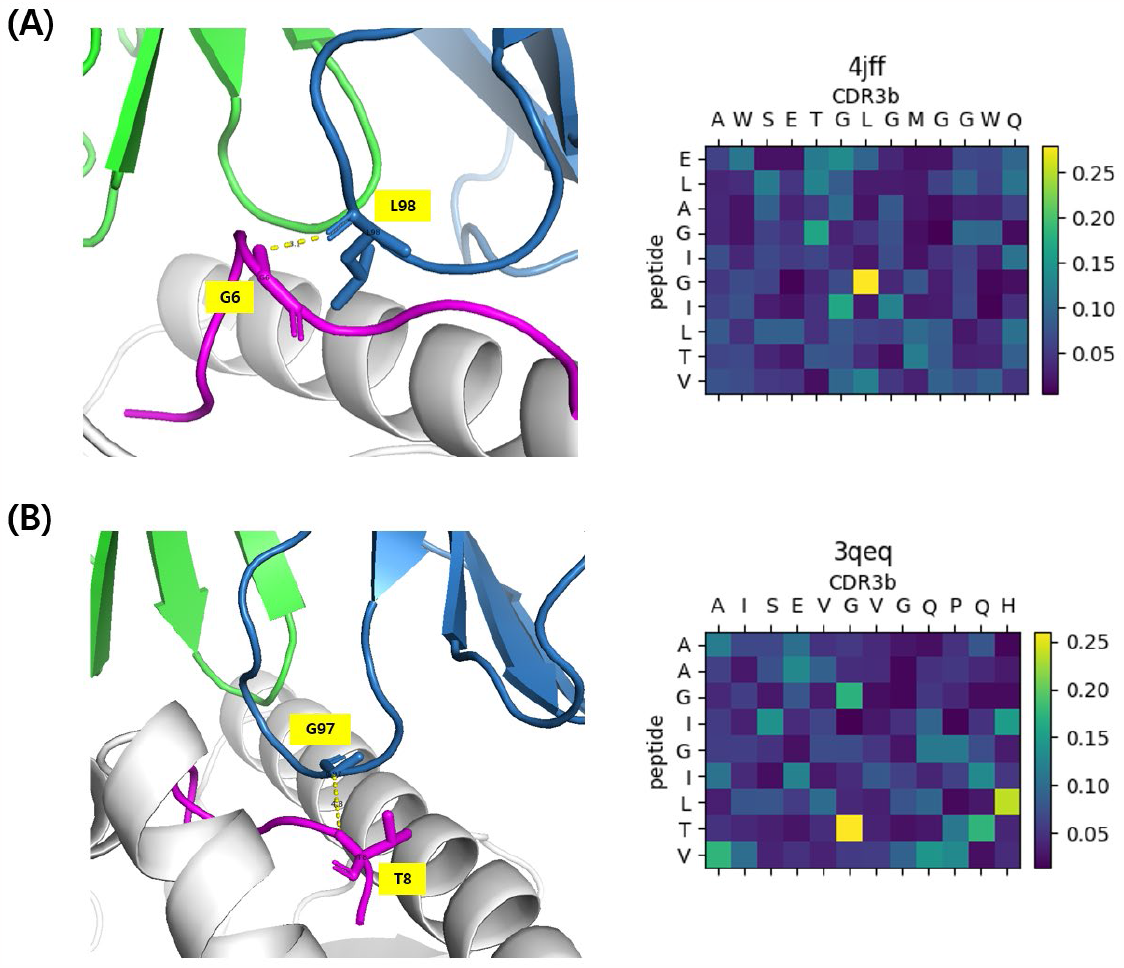
Analysis of crystal TCR-pMHC structures (left) and the attention maps from the reciprocal attention layer (right). The colors in the attention map indicate the attention score. (A) Analysis of the attention map for the structure with PDB code 4JFF. (B) Analysis of the attention map for the structure with PDB code 3QEQ. In the 3D structures, TCR alpha chain is shown in green, TCR beta chain is shown in blue, and the peptide is shown in magenta.

## DISCUSSION

In this work, we propose TSpred, a new ensemble framework combining a CNN architecture with an attention mechanism for the prediction of TCR binding specificity. We take advantage of the unique strengths of each model: the strong ability of CNNs in feature extraction and pattern recognition, and the ability of the attention-based model to focus on more important regions of the input. By formulating an integrated ensemble network, we are able to construct a reliable pan-specific prediction method by harnessing the predictive power of both models in learning TCR-epitope interactions. Based on a comprehensive and rigorous assessment of our model, we find that our model achieves state-of-the-art performances on the prediction task for seen epitopes as well as unseen epitopes. This is a meaningful achievement since most practical applications, such as the development of neoantigen-based vaccines, are related to the prediction of TCR specificity for unseen epitopes (9). Also, in our analysis on the seen epitope datasets, we observe that using paired chain data leads to a significant increase in model performance compared to using beta chain alone, confirming the results from the IMMREP benchmark study (20). In addition, our model shows robust performances across the two widely used negative sampling strategies, while avoiding the bias stemming from each of the strategies. Furthermore, one of the longstanding challenges in this field is the bias caused by peptide imbalance in the training data (7). Although it is a critical factor that affects all predictive methods including ours, TSpred demonstrates the most robust results among all compared predictors when assessed on a balanced dataset. Also, the reciprocal attention mechanism offers model interpretability by showing which residues are key to the interactions of two binding partners. Analysis of the attention maps generated by the model gives us insight into which residue pairs are structurally interacting and thus important to TCR-epitope binding.

With the amount of currently available data, we believe that we have almost reached the limit in increasing the performances of sequence-based TCR binding specificity prediction methods. Despite much effort, model performances for unseen epitopes are still unsatisfactory for use in real clinical applications. In order to increase the model’s reliability in the future, we will need a greater amount of high-quality paired chain data. Current single-cell TCR sequencing data are reported to contain a considerable amount of noise (27). We expect that the development of methods such as ITRAP (27) for denoising single-cell TCR sequencing data will be important in the future. In addition, there are many prediction tools developed in this field, using different types of data, different negative sampling strategies, and different training and testing strategies. As pointed out in the IMMREP benchmark study (20), there is a need for an independent and rigorous benchmark for a thorough evaluation of different methods.

Another possible direction of the research in this field is the development of models based on structural information. Currently, experimental structures of TCR-pMHC complexes are still lacking. However, the latest version of AlphaFold (28) has reportedly achieved a considerable progress in the structure prediction of antibody-antigen complexes, which share many similarities with TCR-pMHC complexes. We anticipate that such progress will lead to better predictions of TCR-pMHC structures, which will facilitate the development of reliable structure-based methods for the prediction of TCR binding specificity.

## DATA AVAILABILITY

Data and source code are available at https://github.com/ha01994/TSpred.

## ACKNOWLEDGEMENTS

We thank Sungjin Choi and the members of the Bioinformatics and Computational Biology Lab for providing helpful advice.

