## Supplementary file for "TSpred: a robust prediction framework for TCR-epitope interactions based on an ensemble deep learning approach using paired chain TCR sequence data"

**Supplementary Fig S1. Modified nested five-fold cross validation.** Unlike standard nested five-fold cross validation, which performs four folds in the inner loop and five folds in the outer loop, we simply use one fold in the inner loop and five folds in the outer loop, as shown below. In the inner loop, hyperparameter tuning is performed using the validation set, and in the outer loop, model evaluation is performed on the test set.

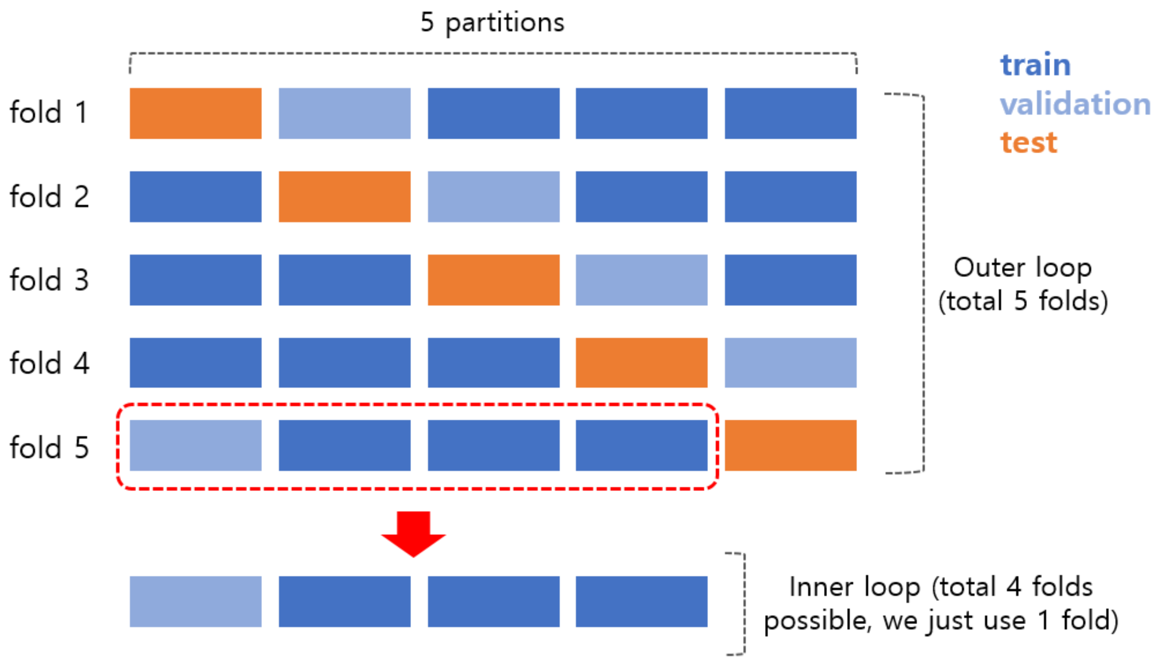

**Supplementary Note S1. Model details and training hyperparameters**

The maximum sequence lengths of CDRα1, CDRα2, CDRα3, CDRβ1, CDRβ2, CDRβ3 are 7, 8, 22 and 6, 7, 23 respectively. The maximum peptide sequence length is 12. The one-hot encoding of the input sequences has a dimension of 21, including 20 amino acids and 1 for padding.

In the CNN-based model, the 1D convolutional layer has an input channel size of 21, an output channel size of 32, a kernel size of 2, and a stride of 1. The max pooling layer has a kernel size of 2 and a stride of 1. The fully-connected layer after the pooling layer outputs vectors of dimension 64. In the attention-based model, the sequence embedding layer has an output dimension of 128. All attention layers are composed of a multi-head attention followed by dropout, residual connection, and layer normalization. They have the following parameters: n_heads=4, d_model=128, d_key=32, and d_value=32.

For both CNN- and attention-based models, we use a batch size of 128. For the CNN-based model, we use a learning rate of 1e-3 and a dropout rate of 0.6, and train for 200 epochs. For the attention-based model, we use a learning rate of 1e-4 and a dropout rate of 0.6, and train for 200 epochs.

### Supplementary Figure S2. Classification results on the NetTCR_full dataset

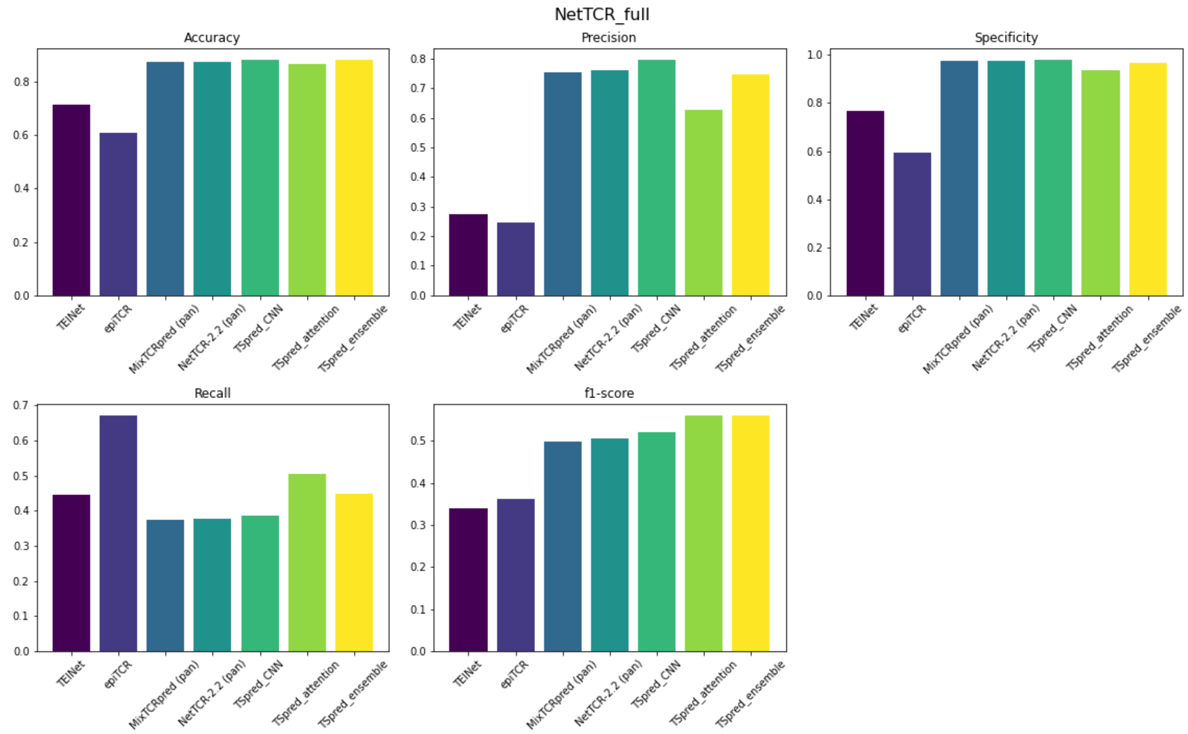

### Supplementary Figure S3. Classification results on the IMMREP dataset

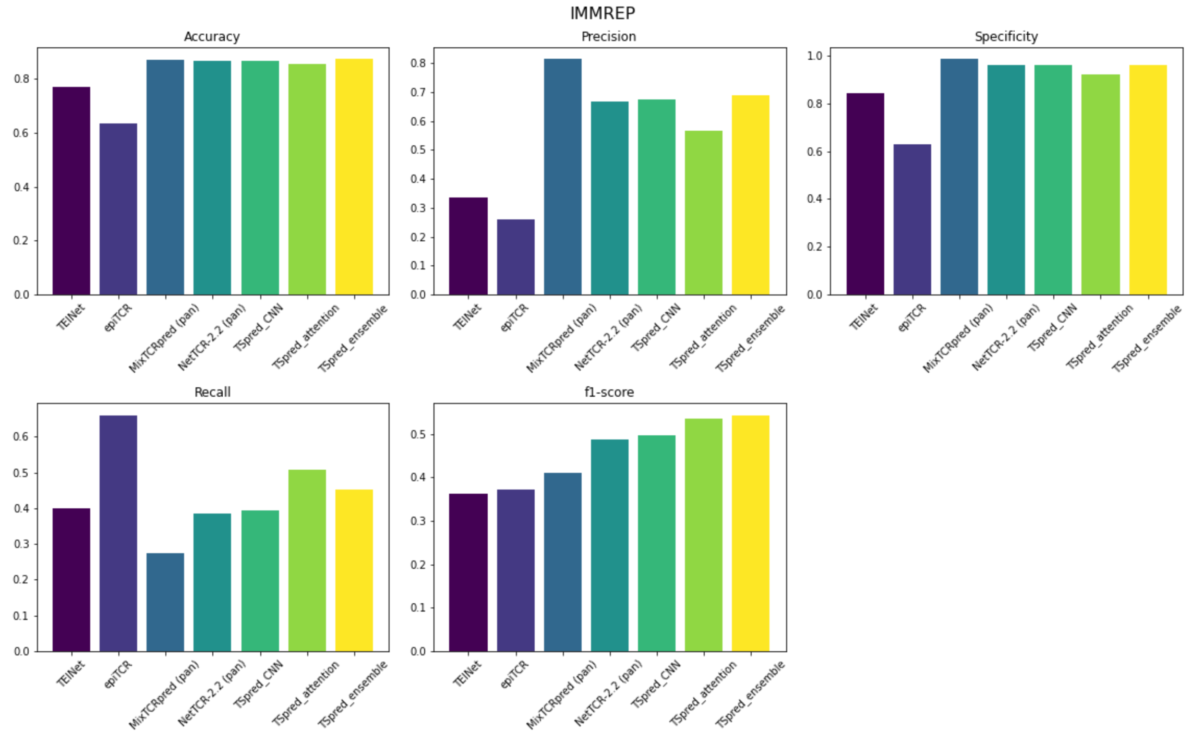

**Supplementary Figure S4. Classification results on the NetTCR_bal dataset**

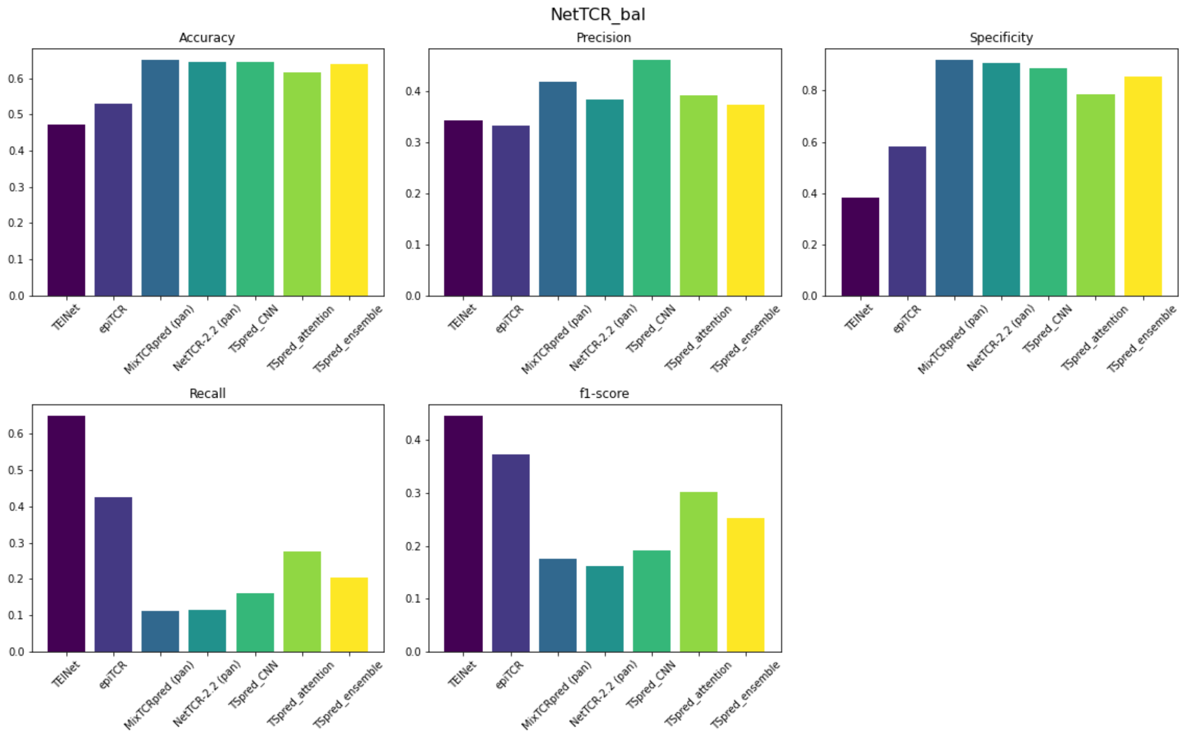

### Supplementary Figure S5. Classification results on the NetTCR_strict dataset

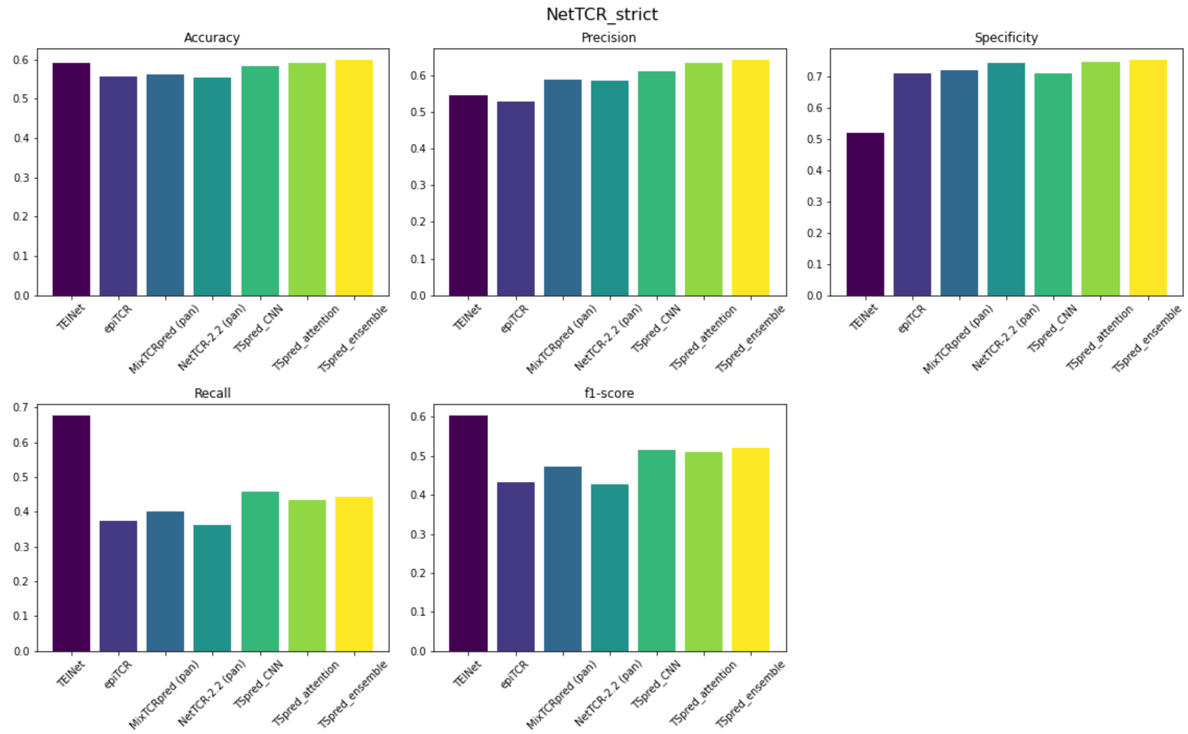
